# Long-read analysis of tetrameric microsatellites with vmwhere supports GGAA repeat length–dependent chromatin state association in Ewing sarcoma

**DOI:** 10.64898/2026.04.08.717017

**Authors:** Sara K. Peterson, A. McCauley Massie, Alexander Rubinsteyn, Jeremy R. Wang, Ian J. Davis

## Abstract

Microsatellites are abundant genomic elements that contribute to genetic diversity and disease-associated regulatory variation. Although long-read sequencing enables accurate resolution of repetitive regions, computational methods for fully resolved microsatellite genotyping remain limited. Here, we introduce variant motif where (vmwhere), a computational framework for identifying, genotyping, decomposing, and visualizing complex tetrameric microsatellites from long-read sequencing data. Using simulated error-free reads, vmwhere accurately measures several genotyping metrics, including allele length, repeat length, maximum consecutive repeat length, and motif density. Applied to long-read whole-genome sequencing data, vmwhere identified sequence interruptions, motif-specific differences in repeat architecture, and ancestry-associated allele variation, including long repeat alleles that exceed short-read sequencing limitations. We applied vmwhere to GGAA microsatellites in Ewing sarcoma, an aggressive pediatric cancer driven by EWS-FLI1 fusion oncoprotein, which binds to microsatellites and remodels chromatin. Genome-wide integration of long-read-defined microsatellite architecture with chromatin accessibility and EWS-FLI1 binding revealed that GGAA repeat structure was associated with chromatin state, with longer consecutive repeat microsatellites exhibiting increased EWS-FLI1 binding and chromatin accessibility. Cell line–specific expansions and contractions of GGAA microsatellite repeat length were associated with gains and losses of chromatin accessibility. Further, we identified haplotype-specific chromatin states, with preferential binding and accessibility at longer alleles. Together, these results establish vmwhere as a scalable framework for resolving population-level microsatellite variation and linking repeat architecture to chromatin state. Repeat structure and length characteristics provides insights into genotype–function relationships at microsatellite repeats in cancer.

## Introduction

Microsatellites are tandemly repeated DNA motifs 1–6 bp in length that are abundant and widely dispersed throughout the human genome. Variation in microsatellite length is a major source of genetic diversity and structural variation, and expansions at specific loci have been implicated in numerous human diseases (Gymrek et al. 2016; Mirkin 2007). Microsatellites are highly polymorphic among human populations, with mutation rates exceeding those of standard nucleotide substitutions (Ellegren 2000; Willems et al. 2016) and common allele frequency differences across ancestries. To date, population-scale genetic studies have largely relied on short-read sequencing, which limits analysis to shorter (< 80 bp) and uninterrupted microsatellite repeats (Ichikawa et al. 2023; Ziaei Jam et al. 2023). As a result, complex repeat structures, including motif interruptions, sequence variants, and longer repeat alleles, remain under characterized, constraining our understanding of how microsatellite variation contributes to population diversity, disease susceptibility, and chromatin states

The release of the complete Telomere-to-Telomere (T2T) reference genome has markedly improved the annotation of repetitive regions, enabling more accurate mapping of expected repeat coordinates and generalized internal structures (Aganezov et al. 2022; Nurk et al. 2022). Although many existing microsatellite genotyping algorithms were originally developed for short-read sequencing (Chiu et al. 2021; Dashnow et al. 2018, 2022; Dolzhenko et al. 2019, 2020; Gymrek et al. 2012; Mousavi et al. 2019; Willems et al. 2017), the emergence of long-read sequencing technologies has opened new opportunities for repeat analysis. Long-read sequencing overcomes key limitations of short-read approaches, including read-length constraints and mapping ambiguity, and is PCR-free, reducing technical artifacts such as stutter (Gymrek 2016). Continued improvements in base-calling accuracy offer increasingly reliable sequence-resolved characterization of complex repeats using long-read data (Mahmoud, Agustinho, and Sedlazeck 2025). These advances have created a need for specialized computational frameworks capable of handling long, interrupted, and highly polymorphic microsatellite alleles. Recent tools such as TRGT, LongTR, ATaRVa, and Straglr (Chiu et al. 2021; Dolzhenko et al. 2024; Sivakumar et al. 2025; Ziaei Jam et al. 2024) have begun to address some of these algorithmic gaps, yet opportunity remains for improved resolution of repeat structure and allelic diversity.

Ewing sarcoma (EwS) offers a compelling biological context in which a specific microsatellite has been linked to chromatin state and gene regulation. EwS is a pediatric bone and soft tissue malignancy driven by the EWS-FLI1 fusion oncoprotein (Delattre et al. 1992). Through the FLI1-derived DNA binding domain, the oncoprotein is directed to canonical ETS DNA binding sites containing a central GGAA sequence. However, the oncoprotein gains neomorphic activity and is also directed to GGAA containing microsatellites, where it remodels chromatin and creates de novo regulatory elements (Gangwal et al. 2008; M. Patel et al. 2012; Riggi et al. 2014). Limited studies at loci near EwS-associated genes have shown that sequence interruptions and repeat architecture, rather than repeat count alone, can alter EWS-FLI1 binding and transcriptional activity. At the 6p25 susceptibility locus, targeted long-read sequencing revealed extensive polymorphism in GGAA repeat length across populations, with both repeat length of both the primary and interrupting sequence correlating with EwS risk (O. W. Lee et al. 2023). Similarly, a microsatellite near *EGR2* showed that single-nucleotide substitutions can alter repeat architecture, ultimately modulating EWS-FLI1 binding and transcriptional output (Grünewald et al. 2015). Difference in GGAA repeat length between populations and between tumor and unmatched normal samples have also been reported at the *NR0B1* locus (Monument et al. 2014). Together, these locus-specific studies highlight the cancer association and regulatory relevance of GGAA microsatellite architecture. However, they have yet to explore whether such relationships extend genome-wide, nor how allele-specific variation in repeat architecture relates to chromatin state across the EwS genome. A key limitation of prior genome-scale analyses is their reliance on reference-derived repeat length annotations, which do not capture population- or sample-specific variation in repeat structure. Consequently, systematic genome-wide analyses integrating read-defined microsatellite architecture with chromatin accessibility and transcription factor binding in EwS remain limited.

To explore these relationships, we developed variant motif where (vmwhere), a computational framework for identification, sequence-resolved genotyping, and decomposition of complex tetrameric microsatellites from long-read sequencing data. We benchmark vmwhere against existing tools using simulated long-read data and apply it to the top 10 most frequent tetrameric repeat motifs, spanning 176,778 loci in 100 genetically diverse individuals from the 1000 Genomes Project (Gustafson et al. 2024), to characterize population-level variation in repeat length and structure. We then focus specifically on GGAA microsatellites in EwS cell lines, integrating repeat length and sequence architecture with chromatin accessibility and EWS-FLI1 binding. Together, this work establishes a scalable framework for resolving population-scale microsatellite variation and demonstrates how sequence-resolved repeat architecture at GGAA microsatellites influences chromatin state in EwS.

## Results

### Computational pipeline for tetrameric microsatellite identification and genotyping from long-read sequencing

We developed variant motif where (vmwhere) as a modular pipeline to identify, genotype, and visualize tetrameric microsatellites from long-read sequencing data (Figure 1). vmwhere pipeline (Figure 1A) comprises three core modules, find, genotype, and visualize, that together enable sequence-resolved characterization of microsatellite alleles at genome scale.

**Figure 1.**
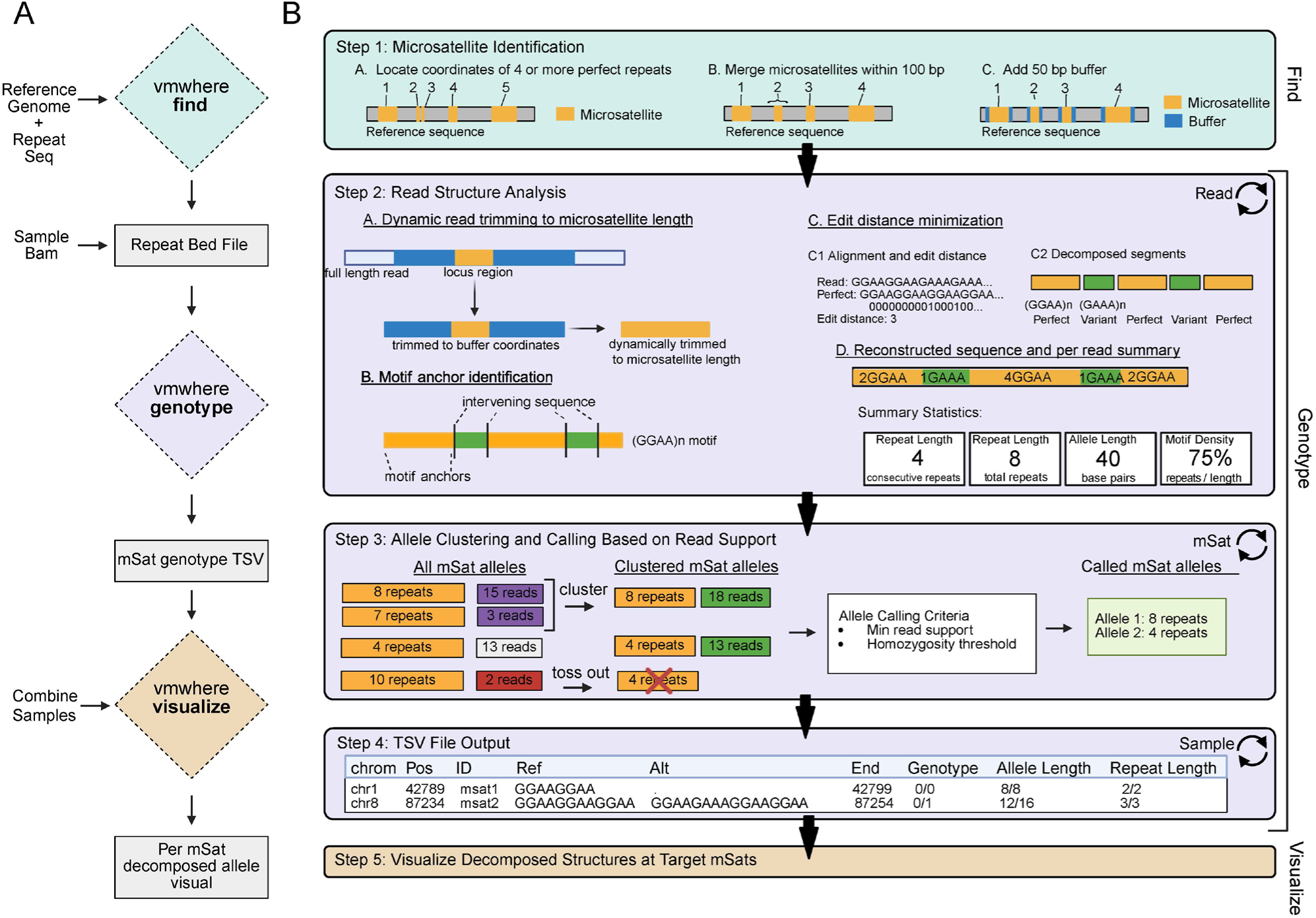
vmwhere pipeline for identification, genotyping, and decomposition of complex tetrameric microsatellites from long-read data. (A) Overview of the vmwhere pipeline, showing required inputs and generated outputs for each module (find, genotype, and visualize). The find module identifies candidate tetrameric microsatellite coordinates from a reference genome sequence, the genotype module performed sequence-resolved allele calling from long-read alignments, and the visualize module generated locus-level summaries across samples. (B) Schematic of the core algorithmic workflow implemented in the genotype module. Reads overlapping candidate loci were filtered and trimmed, canonical motif anchors were identified, and allele sequences were decomposed into motif-consistent and interrupting segments. Reads were clustered into alleles based on edit distance, enabling sequence-resolved allele calling and per-locus visualization across multiple samples.

The find module defines candidate tetrameric microsatellite loci coordinates from a reference genome sequence for a user-specified 4-mer motif (Figure 1B). Tandem repeats exceeding a minimum repeat threshold are detected, nearby occurrences are merged into a single locus based on a user-defined distance, and extended genomic coordinates (tandem repeat start/stop coordinates +/− additional buffer/flanking sequence before and after repeat) are output for downstream genotyping and integration with functional genomics data. Importantly, the reference is used to define the candidate coordinates for genotyping analysis, not for repeat length annotations.

The genotype module performs sequence-resolved allele calling from aligned long reads overlapping each user provided locus. Briefly, this module performs several key steps (Figure 1B). Reads are filtered to retain primary alignments, trimmed to the microsatellite and flanking sequence (based on provided bed file coordinates), and dynamically refined to identify the motif-rich core of each allele. Each allele is decomposed into motif-consistent and interrupting sequence segments, enabling quantification of repeat length, maximum consecutive repeats, motif density, and internal sequence structure. Alleles are called by clustering reads with similar motif architectures, without imposing diploidy assumptions, allowing detection of homozygous, heterozygous, and multi-allelic loci.

Together, vmwhere captures both repeat length variation and motif-level sequence heterogeneity, producing per-sample, per-locus allele summaries suitable for population-scale analysis and integration with epigenomic data. Detailed algorithmic descriptions are provided in the Methods.

### Benchmarking vmwhere against existing long-read genotyping tools on simulated data

To assess genotyping accuracy, we benchmarked vmwhere against four open-source long-read repeat genotyping tools, Straglr (Chiu et al. 2021), LongTR (Ziaei Jam et al. 2024), ATaRVa (Sivakumar et al. 2025), and TRGT (Dolzhenko et al. 2024), using simulated, error-free long-read data generated from 30 tetrameric microsatellite loci present in the reference genome (Figure 2). Benchmark inclusion criteria were tools that are actively maintained, infer repeat length directly from base called read sequences, operate on reference-defined genomic coordinates, and do not require haplotype-resolved or phased BAM inputs. Accuracy was evaluated against simulated truth for multiple genotyping metrics, including allele length, repeat length, maximum consecutive repeat length, and motif density (Figure 2A).

**Figure 2.**
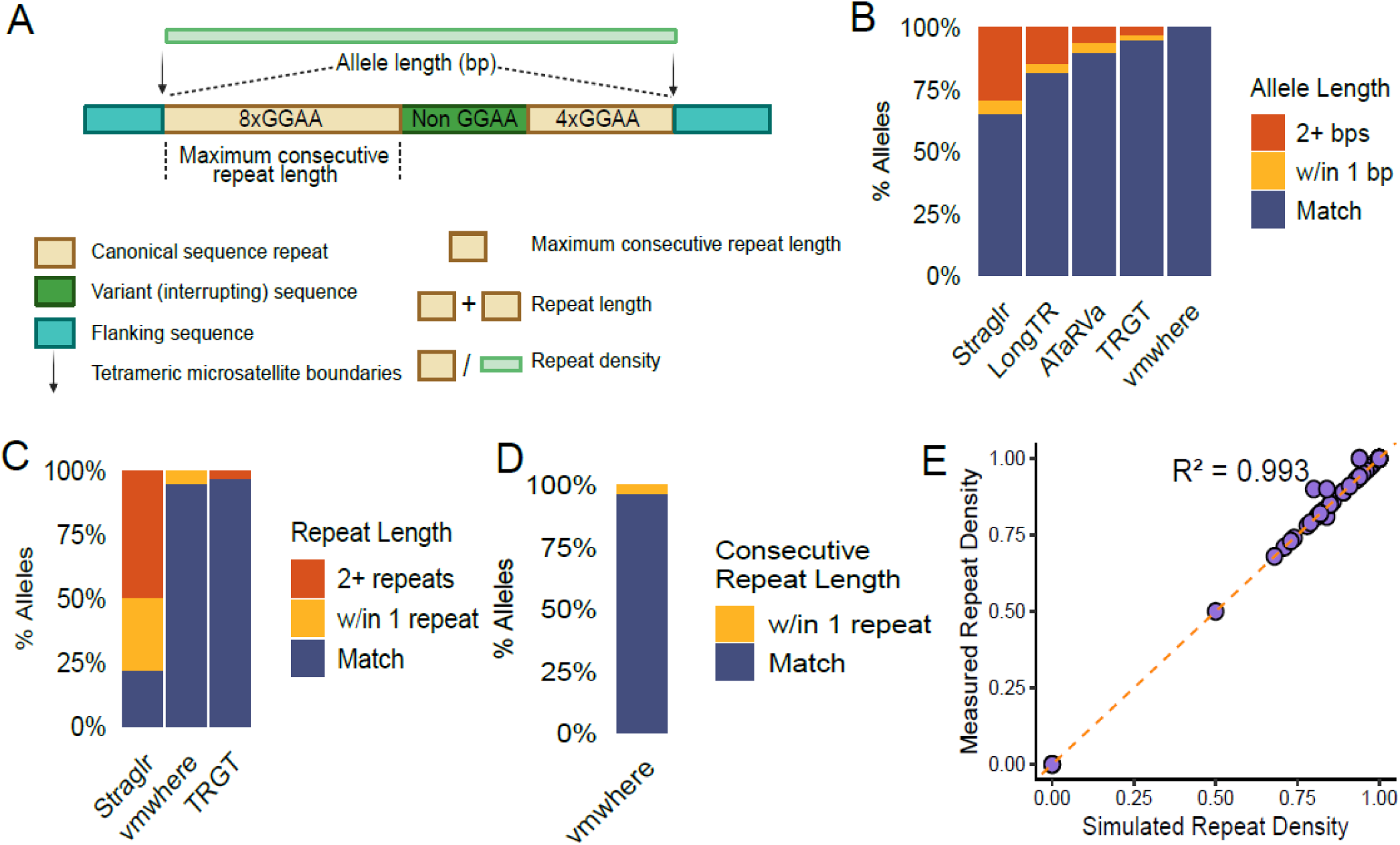
vmwhere accurately genotypes microsatellites from simulated long-read data. (A) Definitions of microsatellite genotyping metrics evaluated, including allele length, repeat length, maximum consecutive repeat length, and motif density. (B) Allele length accuracy, measured as the difference in base pairs between inferred and simulated allele lengths, for Straglr, LongTR, TRGT, ATaRVa, and vmwhere. (C) Repeat length accuracy, measured as the difference in repeat copy number relative to simulated truth, for Straglr, TRGT, and vmwhere. (D) Accuracy of maximum consecutive repeat length estimation for vmwhere, measured as the difference in repeat copy number relative to simulated truth. (E) Comparison of simulated motif density versus measured motif density for vmwhere, demonstrating concordance between inferred and true sequence composition.

Genotyping sensitivity was 100% for vmwhere, TRGT, and ATaRVa, and 97% for LongTR and Straglr. vmwhere achieved the highest accuracy for exact allele length (100%), outperforming TRGT (95%), ATaRVa (90%), LongTR (81,7%), and Straglr (65%) (Figure 2B). With a ±1 bp tolerance, TRGT reached 96.6% accuracy, and ATaRVa reached 93.3%. Notably, vmwhere, ATaRVa, and LongTR correctly resolved allele-specific repeat deletions in non-reference alleles, whereas TRGT and Straglr defaulted to homozygous reference calls, and may therefore underestimate deletions rates at microsatellites.

For repeat length estimation, TRGT showed the highest exact match accuracy (96.7%), followed by vmwhere (95%) and Straglr (21.7%) (Figure 2C). Though TRGT and vmwhere offer similar accuracy for repeat length estimation, only vmwhere detects deletions of repeat sequences. Straglr’s reduced performance could be explained by its reliance on Tandem Repeats Finder, which is optimized for uninterrupted repeats and exhibits lower performance on alleles containing frequent sequence interruptions. TRGT was developed for PacBio HiFi data and embeds error-aware models optimized for HiFi read characteristics. As a result, performance in this benchmark may differ from that reported in the original study, which evaluated TRGT on empirical PacBio datasets rather than idealized, error-free simulations.

vmwhere uniquely reports maximum consecutive repeat length and motif density, metrics derived directly from sequence decomposition. Compared to simulation truth, vmwhere identified maximum consecutive repeat number with 96.7% accuracy (Figure 2D), and motif density showed near-perfect concordance with ground truth (R² = 0.993) (Figure 2E). These features enable discrimination between pure and interrupted alleles and are not accessible through existing genotyping tools.

### Genome-wide identification of tetrameric microsatellites in the T2T human reference

Using vmwhere find, we systematically catalogued tetrameric microsatellite loci in the T2T-CHM13v2 human reference genome. We first defined a set of 42 unique tetrameric microsatellite sequences according to four criteria: (a) non-homopolymeric (e.g., AAAA excluded); (b) combining of reverse complements (e.g., GGAA and TTCC considered equivalent); (c) exclusion of dinucleotide repeats (e.g., ATAT); and (d) exclusion of phased permutations of the same sequence (e.g., GGAA distinct from GAAG, AGGA, and AAGG). vmwhere find applied to the T2T reference genome identified 176,778 tetrameric microsatellite loci with at least four perfect repeats, with repeats within 100 bp merged into a single microsatellite. Per the reference sequence, loci ranged from 4 to 60 maximum consecutive repeat length (4 - 228 total repeat length).

To benchmark sensitivity, we compared vmwhere identified repeat microsatellite loci to those defined by Tandem Repeats Finder (TRF) (Benson 1999) and RepeatMasker (RM) (Tarailo-Graovac and Chen 2009). vmwhere identified 585 tetrameric repeat loci that were not detected by the other tools (Supplementary Figure 1A). All vmwhere uniquely identified tetrameric repeat loci were short (4-5) reference defined repeats (Supplementary Figure 1B), suggesting that statistical thresholds used by TRF and RM may not detect shorter tetrameric microsatellite repeat loci.

Consistent with prior analysis of other human reference genomes (Subramanian, Mishra, and Singh 2003), A-rich motifs (AAAT, AAAC, AAAG) were the most common, followed by G-rich GGAA and AGAT (Supplementary Figure 1C). Whereas most motifs were distributed across both intergenic and intragenic regions, subsets showed bias (e.g., ATCG and ACGT confined to intragenic regions; ACAG, AGCC, CTGG, and AAACT enriched in intergenic regions).

For downstream population-scale analyses, we restricted subsequent analyses to the 10 most frequent tetrameric repeats, which occurred at 154,468 loci, accounting for 87.4% of all tetrameric microsatellite loci in the T2T genome.

### Tetrameric microsatellites exhibit widespread sequence interruptions and variable motif density

First, we applied vmwhere at the previously annotated EwS-associated, highly polymorphic, 6p25 GGAA microsatellite loci (Machiela et al. 2018). vmwhere identified 18 distinct alleles across the cohort, all of which were previously observed (O. W. Lee et al. 2023) and resolved internal sequence interruptions within longer GGAA repeat segments (Figure 3A). The three most common alleles matched prior reports, validating both repeat length estimates and internal sequence decomposition genotyping capabilities of vmwhere on real world long-read data.

**Figure 3.**
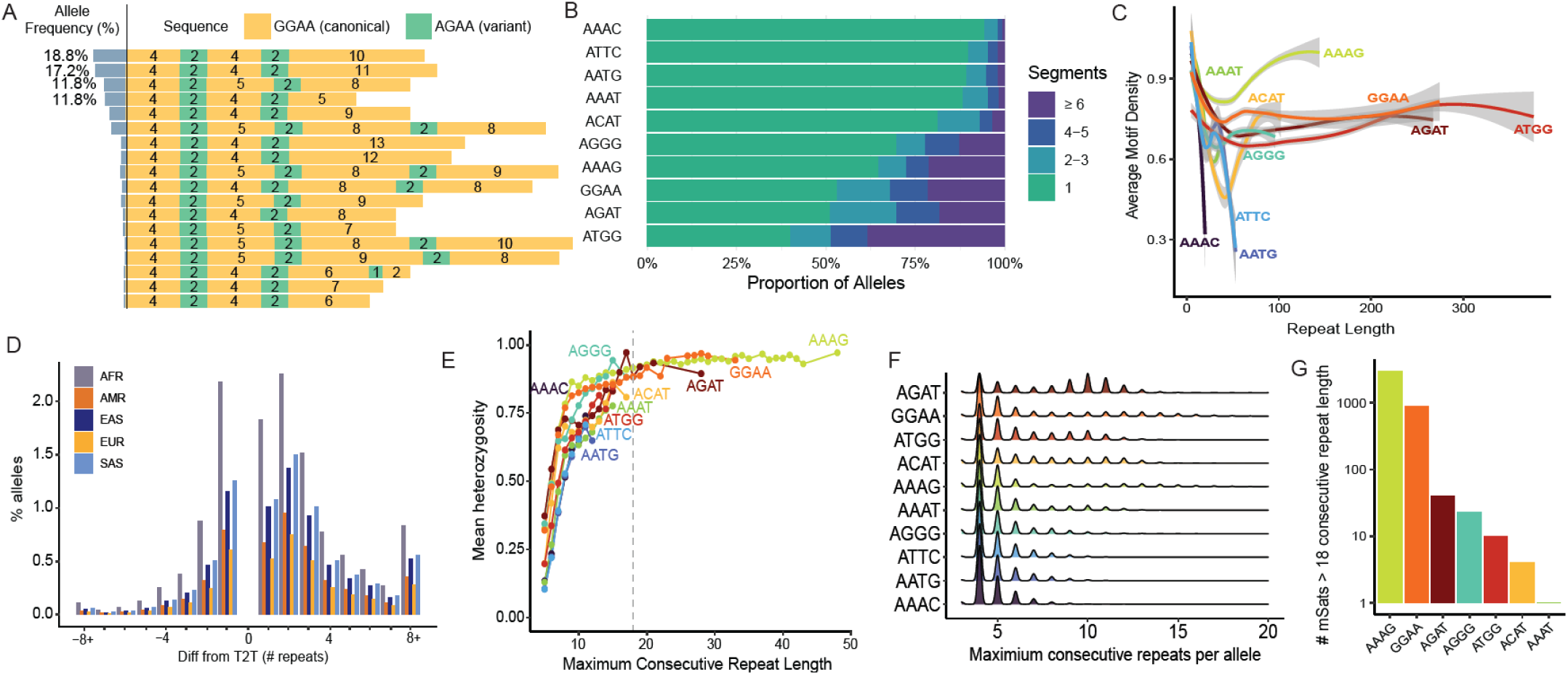
Population analysis of long-read data with vmwhere reveals motif-specific tetrameric microsatellite architectures. (A) Sequence-resolved alleles at the 6p25 Ewing sarcoma susceptibility locus across population samples, with the most prevalent allele frequency indicated. Grey bars represent allele frequencies. Distinct allele structures are shown on the right, with the canonical GGAA motif shown in yellow and the interrupting AGAA motif shown in green. (B) Proportion of alleles for each tetrameric motif stratified by the number of sequence segments, where segments are defined by sequence decomposition of the full allele and each segment represents a change in repeat structure. (C) Relationship between total repeat length and average motif density. Shaded regions indicate 95% confidence intervals. (D) Percentage of variant alleles stratified by ancestry and binned by repeat length difference relative to the T2T reference allele length. (E) Relationship between maximum consecutive repeat length and mean population heterozygosity. (F) Distributions of maximum consecutive repeat length by tetrameric motif. (G) Number of microsatellite loci with allelic support for maximum consecutive repeat length greater than 18, stratified by motif sequence.

We next applied vmwhere to whole-genome Oxford Nanopore Technology (ONT) sequencing data from 100 individuals from the 1000 Genomes Project, spanning five global superpopulations (AFR, AMR, EAS, EUR, SAS) (Gustafson et al. 2024). All population-scale analyses were restricted to loci corresponding to the top 10 most frequent tetrameric motifs identified in the reference as noted previously.

Genome-wide decomposition of alleles revealed that many tetrameric microsatellites exhibited discontinuous architectures (e.g., 4AGAT–2GGAT–4AGAT; three segments) rather than contiguous tandem repeats (e.g., 8GGAA; one segment). Segment counts varied by canonical sequence: AAAC repeats were frequently uninterrupted, whereas ATGG, GGAA, and AGAT loci were commonly interrupted, often comprising two to three distinct segments (Figure 3B). An example of a complex microsatellite dominated by imperfect repeat alleles illustrated the structural complexity that sequence-interrupted microsatellites can display within a population (Supplementary Figure 2A). Consistent with this pattern, longer alleles more frequently exhibited discontinuous architectures, reflecting increased sequence interruption with repeat length.

We next analyzed the sequence of microsatellite interruptions, leveraging the sequence decomposition framework to explore genome-wide patterns of repeat interruption on an allele-by-allele basis. For most motifs (six of ten), the most frequent interruption differed from the primary ed tetranucleotide sequence by a single nucleotide, consistent with SNP-driven motif conversion as a potential source of allelic diversity (Table *1*). This pattern was consistent with a prior genome-wide association study that identified a SNP one class of for which the risk allele altered repeat architecture at one microsatellite locus (Grünewald et al. 2015)this finding across microsatellites. For A-rich repeat motifs, AAAT and AAAC, the most common interruption was a single adenine nucleotide; however, these interruptions were observed in a smaller fraction of alleles relative to the dominant interruptions identified for other motifs (Table 1).

**Table 1.** For each tetrameric microsatellite motif, allele frequency, maximum consecutive repeat length, total repeat length, and motif density of the allele exhibiting the longest consecutive repeat were summarized.

| Primary Motif | Interruption Sequence | Allele Prevalence (%) | 250 |
| --- | --- | --- | --- |
| ATGG | GTGG | 52 |  |
| AGAT | GAT | 48 | 251 |
| GGAA | AGAA | 48 |  |
| AAAG | AG | 38 | 252 |
| AGGG | AGGA | 40 | 253 |
| ACAT | ACGT | 34 |  |
| AAAT | A | 24 | 254 |
| AATG | AGTG | 27 | 255 |
| ATTC | ACTC | 27 |  |
| AAAC | A | 23 | 256 |

We next examined how motif density changed as a function of total repeat length across canonical repeat sequences. AAAG motifs maintained the highest motif density with increasing repeat length, whereas AATC and AAAC repeats showed a sharp decline in density as repeat length increased (Figure 3C). These patterns indicated that core sequence continuity was dependent on both repeat motif and allele length.

Overall, these results indicated that interruptions to the core sequence were common across tetrameric microsatellite motifs and frequently affected a substantial proportion of alleles, highlighting interruption structure as a major contributor to allelic variation in microsatellite architecture.

### Sequence-dependent heterozygosity and ancestry-associated allelic variation

We next quantified allelic diversity across the five superpopulations (AFR, AMR, EAS, EUR, SAS). Consistent with prior findings using short-read sequencing in alternate reference genomes (Shi et al. 2023) (e.g., GRCh38), long-read based analyses demonstrated that most alleles matched the reference regardless of ancestry, and variant allele frequency declined with increasing divergence from the reference (Figure 3D). As reported in prior studies (Kinney et al. 2019; Willems et al. 2014), African individuals exhibited the greatest allelic variation, particularly at tetrameric repeats (Figure 3D).

Importantly, this diversity was likely under quantified in short-read analyses due to limited resolution of longer repeats. Using vmwhere, we identified 11,251 tetrameric microsatellite loci with alleles exceeding 72 bp in repeat length. Long alleles comprised 3.2–3.4% of total alleles across superpopulations (Supplementary Figure 2B). Individuals of African ancestry accounted for 31.3% of all long alleles identified (Supplementary Figure 2C).

Because vmwhere uniquely quantified maximum consecutive repeat length, we examined heterozygosity as a function of this feature. Heterozygosity increased with longer consecutive repeat lengths, with GGAA, AAAG, and AGAT loci showing the strongest length-associated effects (Figure 3E). These findings extend prior short-read studies (Payseur, Jing, and Haasl 2011; Shi et al. 2023; Sun et al. 2012; Willems et al. 2014; Ziaei Jam et al. 2023), as allelic support for repeat lengths exceeding short-read resolution limits was observed (i.e., alleles to the right of the gray dashed line on Figure 3E).

### Sequence specific enrichment of longer uninterrupted tetrameric microsatellites

Analysis of maximum consecutive repeat length revealed strong sequence dependence. Some tetrameric microsatellites, such as AAAC, and AATG occurred predominantly as shorter consecutive repeats, while others, AAAG, GGAA, and AGAT exhibited bimodal distributions, with higher frequencies at both shorter and longer consecutive repeat lengths (Figure 3F).

Among loci corresponding to the top 10 motifs, vmwhere identified 3,892 loci (2.5% of all tetrameric microsatellites) with alleles exceeding 18 consecutive repeats. Seven motifs (AAAG, GGAA, AGAT, AGGG, ATGG, ACAT, and AAAT) comprise longer alleles, with AAAG and GGAA accounting for 98% of loci with extended consecutive repeat lengths (Figure 3G). These loci included alleles with extreme lengths that reached up to 144 pure repeats for AAAG (Supplementary Figure 2D) and 68 for GGAA (Supplementary Figure 2E).

Long alleles generally occurred at lower allele frequencies (0.012 – 0.02) and, when present, were predominantly uninterrupted pure microsatellites in 4 of the 6 motifs (AAAT excluded due to single long locus, n = 1) (Table *2*). These observations indicated that extended uninterrupted tetrameric repeats were present in the population despite high mutation rates.

**Table 2.** Repeat length and allele frequency of longest consecutive allele per motif.

| Motif | Allele Frequency | Maximum Consecutive Repeat Length | Total Repeat Length | Motif Density |
| --- | --- | --- | --- | --- |
| AAAG | 0.012 | 144 | 144 | 1 |
| GGAA | 0.018 | 68 | 68 | 1 |
| AGAT | 0.020 | 32 | 36 | 0.94 |
| AGGG | 0.015 | 29 | 29 | 1 |
| ACAT | 0.016 | 26 | 26 | 1 |
| ATGG | 0.015 | 25 | 29 | 0.97 |

Together, these results demonstrated that sequence-resolved analysis of long-read data using vmwhere captured substantial population-level variation in tetrameric microsatellite architecture, including motif-specific differences in repeat purity, length distributions, and heterozygosity. GGAA microsatellites in particular frequently exhibited long maximum consecutive repeat lengths and high population heterozygosity, motivating their focused analysis in disease contexts.

### Maximum consecutive repeat length is associated with chromatin state at GGAA microsatellites in Ewing sarcoma

Because of the functional relationship between microsatellites and Ewing sarcoma, we applied vmwhere to long-read whole genome sequencing from four established EwS cell lines (A673, SK-ES-1, SK-N-MC, MHH-ES-1). We then integrated microsatellite genotypes with EWS-FLI1 binding, chromatin accessibility, and expression of genes within 100 kb of each locus. Across all four cell lines, both EWS-FLI1 binding and chromatin accessibility broadly increased with GGAA microsatellite length (based on the longest detected allele), consistent with prior observations of EWS-FLI1 binding based on GRCh38 reference repeat length annotations (Figure 4A, Supplementary Figure 3A) (Johnson, Taslim, et al. 2017).

**Figure 4.**
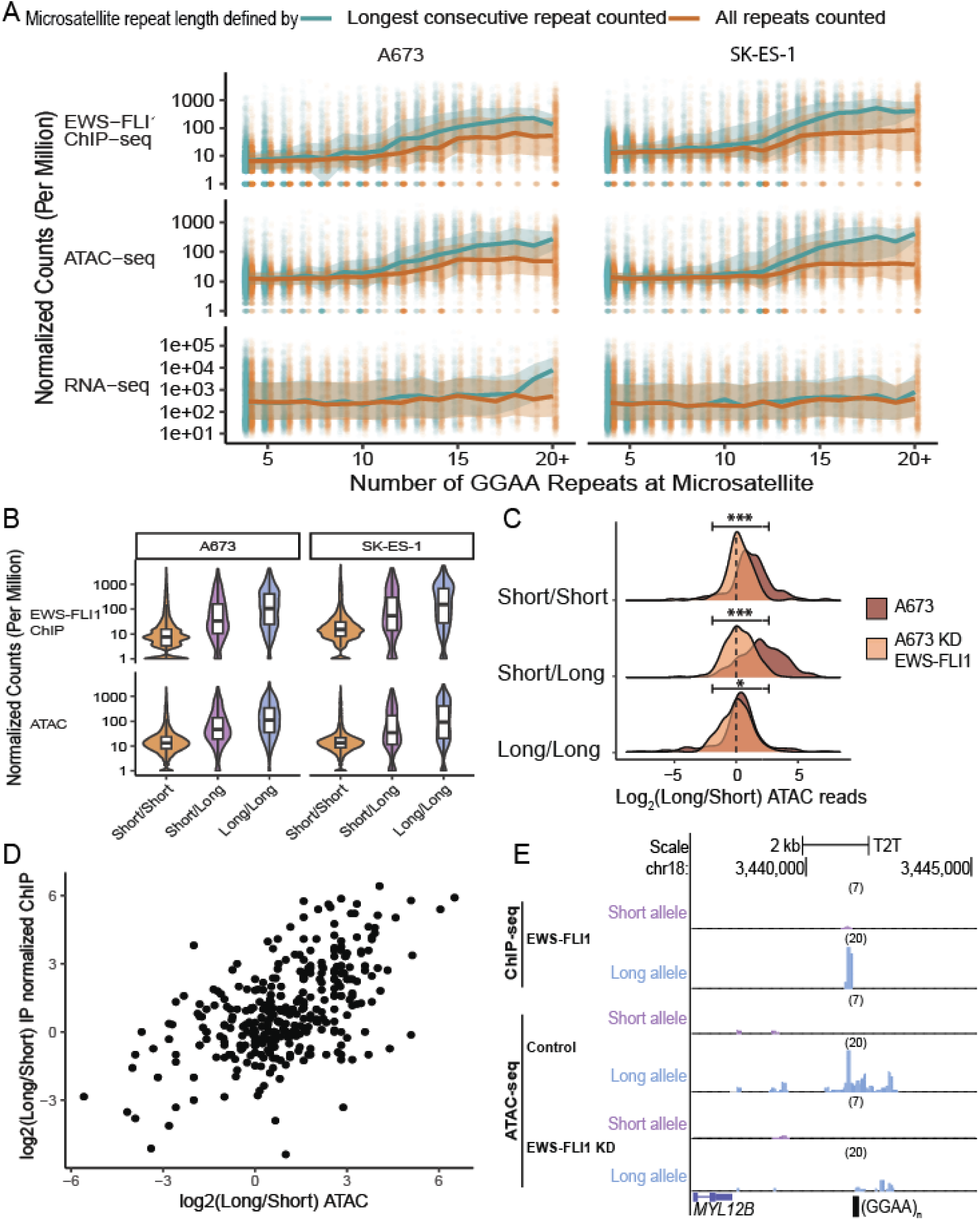
GGAA microsatellite consecutive repeat length and haplotype are associated with chromatin accessibility and EWS-FLI1 binding in Ewing sarcoma. (A) Relationship between GGAA longest allele repeat length per microsatellite, as determined by vmwhere, and normalized signal (counts per million) at the same microsatellite in two Ewing sarcoma cell lines, A673 (left) and SK-ES-1 (right). Shown are EWS-FLI1 ChIP–seq (top), ATAC–seq (middle), and RNA–seq (bottom). Signal is quantified once per sequencing modality per microsatellite and stratified by either maximum consecutive repeat length or total repeat length. Line represents the median signal, with shading showing the interquartile (Q1 and Q3) range. (B) EWS-FLI1 binding and chromatin accessibility at GGAA microsatellites stratified by haplotype, with alleles classified as short (≤ 11) or long (≥ 12) based on maximum consecutive repeat length. (C) Distribution of allele-specific signal ratios (long/short allele) for EWS-FLI1 binding and chromatin accessibility at heterozygous GGAA microsatellites in A-673 cells, shown under control conditions and following EWS-FLI1 knockdown. Statistical significance is assessed using a Wilcoxon rank-sum test (p < 0.05, p < 0.01, p < 0.001). (D) Correlation between allele-specific chromatin accessibility and allele-specific EWS-FLI1 binding at the same GGAA microsatellite loci, shown as log₂ fold change reads aligned to the long allele divided by reads aligned to the shorter allele for ATAC–seq versus ChIP–seq signal (Pearson r = 0.53). (E) Example GGAA microsatellite locus in A673 exhibiting a short/long haplotype (7 versus 20 consecutive repeats). EWS-FLI1 binding and chromatin accessibility preferentially localize to the longer allele, and this allelic preference is reduced following EWS-FLI1 knockdown.

Matched by the same total number of repeats, loci with longer uninterrupted repeat segments exhibited higher accessibility and binding. In contrast, neither total nor consecutive repeat length was associated with overall variation in nearby gene expression. These results were consistent with GGAA microsatellites functioning as regulatory elements for a subset of EwS associated target genes.

Length-dependent accessibility was lost following EWS-FLI1 knockdown in A673 cells (Supplementary Figure 3B), indicating that the observed chromatin effects are mediated by EWS-FLI1. For comparison, no length-associated accessibility or EWS-FLI1 binding was observed at AGAT tetrameric microsatellites (Supplementary Figure 3C), which occurred at similar genomic prevalence and exhibited comparable consecutive repeat length distributions, consistent with known motif specificity for EWS-FLI1. This also indicates that microsatellite associated chromatin accessibility is specific to those that contain GGAA.

To quantify the relationship between GGAA repeat length and chromatin accessibility, we modeled accessibility as a function of maximum consecutive GGAA repeat length using a piecewise linear regression with a hinge at 11 repeats. Below this threshold, accessibility showed a weak dependence on repeat length, increasing modestly with each additional repeat (β = 0.15 log₂ CPM per repeat). In contrast, above 11 repeats accessibility increased sharply and approximately linearly, with a greater than threefold increase in slope (additional β = 0.33; combined slope = 0.50 log₂ CPM per repeat).

Applying the same modeling framework to total repeat length yielded a weaker association. While accessibility showed a modest increase with total repeat length below 11 units (β = 0.12 log₂ CPM per repeat), little additional dependence was observed above this threshold (additional β = 0.03; combined slope = 0.15 log₂ CPM per repeat). Together, these results indicated that maximum consecutive GGAA repeat length was more informative than cumulative repeat length for predicting chromatin accessibility at GGAA microsatellites. Further, they support EWS-FLI1 activity based on uninterrupted repeat sequences, rather than the mix continuous and interrupted motifs.

### Allele-specific chromatin accessibility at GGAA microsatellites

To date, analyses of chromatin accessibility have yet to explore allelic contribution. We hypothesized that chromatin state may be preferentially driven by the longest allele at a microsatellite locus. We classified alleles as short (≤ 11 repeats) or long (> 11 repeats) and stratified microsatellites by haplotype composition. Haplotype composition was defined using the per-locus allele calls generated by vmwhere, with haplotypes assigned based on the combination of allele lengths present at each microsatellite locus. Microsatellites with short/short haplotypes exhibited the lowest levels of EWS-FLI1 binding and chromatin accessibility, whereas short/long loci showed substantially increased normalized signal (Figure 4B, Supplementary Figure 3D). Notably, after EWS-FLI1 knockdown haplotype associated accessibility was decreased at haplotypes which contained one or two long alleles (Supplementary Figure 3E).

We next quantified allele-specific chromatin accessibility in A673 EwS cell line at 501 heterozygous GGAA microsatellite loci by aligning reads to a haplotype-resolved assembly. At long/short loci, chromatin accessibility was preferentially enriched on the longer allele but showed more balanced binding at short/short and long/long loci, with a small preference for the long allele (Figure 4C). This allelic imbalance was markedly reduced following EWS-FLI1 knockdown, accessibility distributions collapsed toward symmetry across all haplotypes. Paired comparisons of allele-specific accessibility between control and knockdown conditions revealed significant shifts in accessibility distributions for each haplotype class (paired Wilcoxon signed-rank test, p < 0.05, p < 0.01, p < 0.001). These results indicated that allele-specific chromatin accessibility was EWS-FLI1 dependent.

Allele-specific EWS-FLI1 binding showed a similar preference for the longer allele at long/short microsatellites (Supplementary Figure 3F), this mirrored chromatin accessibility results. Broadly at GGAA microsatellite loci, allele-specific chromatin accessibility was moderately positively correlated with allele-specific EWS-FLI1 binding (Pearson r = 0.55), indicative of coordinated allele-specific occupancy and chromatin remodeling (Figure 4D). A representative example is the GGAA microsatellite downstream of MYL12B, a gene involved in modulating myosin II activity that is essential for cytoskeletal organization, which exhibited a long/short haplotype (20 vs. 7 consecutive repeats). At this locus, chromatin accessibility and EWS-FLI1 binding were preferentially enriched on the longer allele and chromatin accessibility was reduced following EWS-FLI1 knockdown (Figure 4E).

### Maximum consecutive repeat length expansions and contractions at GGAA microsatellites are associated with differential chromatin accessibility in Ewing sarcoma

We next assessed whether cell line-specific differences in GGAA microsatellite length were associated with variation in chromatin accessibility. First, we identified loci with differential accessibility in each EwS cell line using a one-versus-rest framework. Next, we used the same one-versus-rest framework to classify microsatellites based on maximum consecutive repeat length as expanded, contracted, or within range. Microsatellites were classified and expanded or contracted based on a repeat length difference of ≥ 2 consecutive repeats relative to the largest or smallest value observed across other cell lines. The threshold for expansion and contraction was chosen to reflect the biochemically predicted EWS-FLI1 binding at GGAA microsatellites, in which binding occurs approximately every other repeat unit (Johnson, Mahler, et al. 2017). Expanded and contracted microsatellites together comprised 5-6% of all microsatellites (Figure 5A). Length changes at most expanded and contracted loci differed by fewer than five consecutive repeats relative to min/max lengths from other samples (Figure 5B, Supplementary Figure 4A).

**Figure 5.**
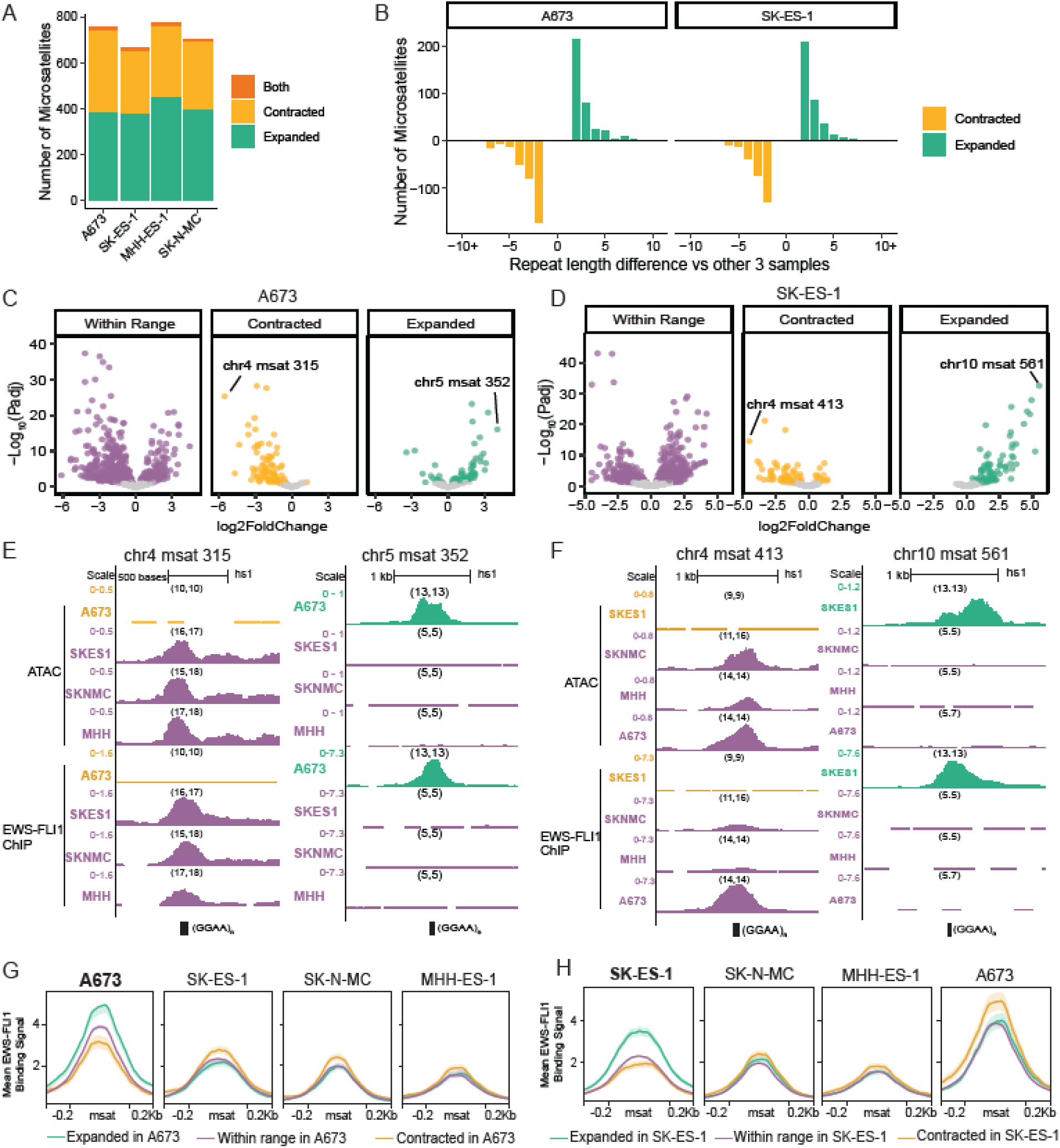
Sample-specific changes in GGAA microsatellite length are associated with coordinated changes in chromatin accessibility and EWS-FLI1 binding. (A) Classification of GGAA microsatellites by repeat length variation in each EwS cell line. Microsatellites are classified as expanded (≥ 2 consecutive repeats longer than the maximum observed in other cell lines), contracted (≤ 2 consecutive repeats shorter than the minimum observed in other cell lines), or within range. A subset of loci shows allelic support for both expansion and contraction within the same cell line. (B) Distribution of consecutive repeat length differences observed among expanded and contracted GGAA microsatellites across cell lines. (C) Volcano plot of differential chromatin accessibility in the A673 cell line, stratified by GGAA microsatellite length change category. The x-axis is log_2_fold change and the y-axis is –log_10_ (adjusted p-value). Microsatellites within range of other cell lines (left), with allelic support for contraction (middle), or with allelic support for expansion (right) are shown. Points with p > 0.05 are shown in grey, and significant loci (p < 0.05) are colored by microsatellite change category. (D) Volcano plot of differential chromatin accessibility in the SK-ES-1 cell line, stratified by GGAA microsatellite length change category, as in panel C. (E) Example GGAA microsatellite loci in A673 showing the largest decrease in chromatin accessibility associated with repeat contraction (left) and the largest increase associated with repeat expansion (right). ATAC–seq (top) and EWS-FLI1 ChIP–seq (bottom) signals are shown across all four EwS cell lines. Numbers in parentheses indicate maximum consecutive repeat lengths for the two alleles in each sample at that locus. (F) Example GGAA microsatellite loci in SK-ES-1 showing the largest decrease in chromatin accessibility associated with repeat contraction (left) and the largest increase associated with repeat expansion (right), displayed as in panel E. Numbers in parentheses indicate maximum consecutive repeat lengths for the two alleles in each sample at that locus. (G) Mean EWS-FLI1 binding signal across GGAA microsatellite loci stratified by expansion status as defined in A673 and shown across all cell lines. (H) Mean EWS-FLI1 binding signal across GGAA microsatellite loci stratified by variation status as defined in SK-ES-1 and shown across all cell lines.

Microsatellites classified as either expanded or contracted exhibited significantly larger absolute fold changes in chromatin accessibility compared to unchanged microsatellites, a trend observed in all cell lines (Supplementary Figure 4B). Moreover, sample-specific expanded microsatellites were frequently associated with increased chromatin accessibility (one sample vs remaining three), whereas contracted microsatellites were associated with decreased accessibility. In contrast, unchanged microsatellites showed a balanced distribution of accessibility increases and decreases. This trend was observed in all cell lines (Figure 5C, D, Supplementary Figure 4C, D). Together, these results demonstrate coordinated gains and losses of chromatin accessibility at loci that have undergone repeat length changes.

Example loci illustrating expansion-associated gains and contraction-associated losses of chromatin accessibility are shown for the microsatellites with the largest significant fold change in each direction for A673 (Figure 5E), SK-ES-1 (Figure 5F), MHH-ES-1 (Supplementary Figure 4C), and SK-N-MC (Supplementary Figure 4D). In A673, a contracted GGAA microsatellite on chromosome 4 exhibited a homozygous maximum consecutive repeat length of 10, whereas the same locus ranged from 15–18 consecutive repeats in the other cell lines. At this locus, chromatin accessibility and EWS-FLI1 binding was reduced in A673 relative to the other cell lines (log₂FC = −5.54). Conversely, an expanded GGAA microsatellite on chromosome 5 in A673 exhibited a homozygous maximum consecutive repeat length of 12, compared to a homozygous length of 5 in the remaining cell lines. At this locus, both chromatin accessibility and EWS-FLI1 binding was increased in A673 relative to the rest (log₂FC = 4.89).

In addition to changes in chromatin accessibility, aggregate EWS-FLI1 binding at the same GGAA microsatellites showed concordant, length-dependent behavior. When microsatellites were stratified by A673-defined expansion status, EWS-FLI1 binding was highest at loci classified as expanded in A673, intermediate at loci within range, and lowest at contracted loci within A673 (Figure 5G). When these same locus sets were examined in the remaining cell lines, A673-defined contracted loci exhibited higher average EWS-FLI1 binding relative to A673-defined expanded or within-range loci, consistent with the longer repeat lengths observed at these loci in the other cell lines. This reciprocal pattern further supported a model in which EWS-FLI1 binding often reflects relative GGAA repeat length across samples. This trend was consistently observed across all cell lines, including SK-ES-1 (Figure 5H), MHH-ES-1 (Supplementary Figure 4E), and SK-N-MC (Supplementary Figure 4F). Taken together, these data suggest that while reference genome-based microsatellite length determination has informed our general appreciation for the association between consecutive repeat length and EWS-FLI1 activity on chromatin, differences across cell lines are likely to result from cell line specific variation in consecutive repeat length.

## Discussion

In this study, we present vmwhere, a comprehensive pipeline to identify, genotype, decompose, and visualize tetrameric microsatellites from long-read sequencing data. Using simulated long-read data, we benchmarked vmwhere against four publicly available genotyping tools and demonstrated high accuracy across multiple metrics, including allele length, repeat length, maximum consecutive repeat length, and motif density. Application of vmwhere to population-scale long-read data further revealed substantial sequence heterozygosity and ancestry-associated allelic variation at tetrameric microsatellite loci, extending prior observations limited by short-read sequencing limitations.

Across the genome, GGAA microsatellites emerged as a distinct class of tetrameric repeats, characterized by bimodal length distributions and enrichment of long, consecutive repeat lengths. Leveraging this sequence-resolved framework in EwS, we identified a genome-wide association between maximum consecutive GGAA repeat length and chromatin state, specifically chromatin accessibility and EWS-FLI1 binding. Accessibility and binding increased above a threshold of approximately 11 consecutive repeats, and allele-specific analyses demonstrated preferential EWS-FLI1 occupancy and chromatin accessibility on the longer allele at heterozygous loci. Finally, we observed a small number of GGAA microsatellites undergoing sample-specific expansions or contractions. These loci were associated with corresponding gains or losses in chromatin accessibility, linking consecutive repeat length variation to changes in chromatin state in EwS.

Long-read sequencing provides an opportunity to study microsatellite variation in ways that were previously inaccessible. Beyond enabling improved genotyping through PCR-free library preparation and the ability to span entire repeat loci within single reads, long-read data support targeted, locus-focused interrogation of microsatellites. Our population-scale analyses indicate that longer and interrupted tetrameric alleles often occur at low frequencies and are therefore likely underrepresented in standard whole-genome datasets. Deep, targeted interrogation of microsatellite loci may be required to capture this variation robustly. Long-read sequencing strategies that support targeted enrichment, such as adaptive sampling on Oxford Nanopore platforms, offer a practical means to achieve the depth necessary to resolve rare alleles and multiallelic loci. Such approaches have the potential to uncover previously overlooked microsatellite variation with functional relevance, including variation that may contribute to disease susceptibility (Miller et al. 2021). Applied at scale across diverse populations, these strategies could enable systematic investigation of how microsatellite architecture shapes genetic diversity. Notably, one limitation of our population-scale variation study is the reliance on the T2T-CHM13v2 reference genome, which represents a single individual. Future incorporation of pangenome reference may better capture ancestry-specific microsatellite diversity.

Our results support the current conceptual shift in the broader tandem repeat field. Rather than representing microsatellites by a single repeat length metric, features such as maximum consecutive repeat length and internal sequence interruptions emerge as distinct and potentially biologically informative properties. Decomposition of read-defined microsatellite alleles into contiguous sequence segments reveals architectural and allele specific variation that is not captured by reference-derived length annotations. This perspective has important implications for genome-wide association studies, in which disease-associated single-nucleotide variants frequently overlap complex and discontinuous repeat loci and may alter repeat architecture rather than acting independently (Grünewald et al. 2015; Postel-Vinay et al. 2012). More broadly, studies of population-scale tandem repeat variation have demonstrated that uninterrupted repeat length of the primary sequence and length of sequence interruptions can be also be predictive of pathogenic potential or disease susceptibility, in addition to total repeat length (J.-M. Lee et al. 2019; O. W. Lee et al. 2023). Together, these observations motivate a view of microsatellites as sequence elements whose structure and composition contribute to genetic variation and may provide new insight into pathogenicity.

Sample-to-sample variation in GGAA microsatellite architecture provides a potential mechanism for explaining a portion of inter-patient heterogeneity in EwS. GGAA microsatellites targeted by the EWS-FLI1 fusion oncoprotein function as regulatory elements and repeat architecture influences chromatin accessibility and transcription factor binding (Gangwal et al. 2008; Riggi et al. 2014). The observation that modest changes in GGAA repeat length are associated with measurable differences in chromatin accessibility and EWS-FLI1 binding suggests that large-scale microsatellite instability is not required for functional effects on chromatin state. Instead, EwS appears sensitive to small architectural changes at a subset of microsatellites, consistent with prior locus-specific studies demonstrating that subtle alterations in GGAA repeat structure can modulate EWS-FLI1 binding and transcriptional output (Grünewald et al. 2015; Postel-Vinay et al. 2012). These studies have proposed a germline–somatic co-option model, in which inherited GGAA repeat variation contributes to disease susceptibility (O. W. Lee et al. 2023; Monument et al. 2014; Postel-Vinay et al. 2012). Extending this framework genome-wide, our results suggest that germline-defined microsatellite architecture may influence chromatin state in established tumors, providing a potential source of some heterogeneity across patients. One further limitation is allele-specific analyses were restricted to heterozygous loci with sufficient uniquely mappable reads, limiting the number of loci that could be interrogated. Emerging long-read–based epigenomic approaches, such as Fiber-seq, may enable more comprehensive characterization of allele-specific chromatin states. These emerging methods, used in connection with vmwhere, would provide broader allele specific analysis at microsatellites in future studies. Taken together, this works represents an important next step toward understanding how microsatellite variation is associated with chromatin state in EwS.

## Methods

### vmwhere algorithm and implementation

vmwhere is freely available as a Python package via PyPl or at GitHub (https://github.com/pirl-unc/vmwhere). The software is implemented in Python and R and provides a command-line interface with three subcommands: find, genotype, and visualize. Installation instructions and usage guidelines are available in the GitHub repository documentation.

### find module

The vmwhere find module identifies microsatellite loci by performing an exact string-based search for a user-specified motif within a reference FASTA sequence. For each contig, the algorithm scans the reference sequence for exact matches to the motif on both the forward strand and its reverse complement. Each detected motif match serves as a seed position from which the algorithm extends sequentially in motif-length increments to identify consecutive, perfectly matched repeat units.

Extension terminates when the next motif-length window no longer matches the motif sequence. The start and end coordinates of the maximal contiguous repeat block are recorded, along with the total number of repeat units detected. Only exact (perfect) motif matches are counted on this first pass of the algorithm.

After scanning the full reference, repeat loci containing fewer than a user-defined minimum number of perfect repeats are dropped. Remaining repeat loci are consolidated through a merging process that combines loci whose coordinates fall within a specific maximum gap distance, producing a single consolidated microsatellite region. Following merging, a user-defined buffer is added upstream and downstream of each merged locus to capture local flanking sequence context. Final merged regions are assigned unique identifiers and reported as BED-formatted coordinates representing buffered microsatellite loci.

### genotype module

The vmwhere genotype module performs microsatellite genotyping by analyzing sequencing reads aligned to user-specified loci and characterizing the structural composition of each allele. User input consists of BED-formatted coordinates (typically output from the find module), an aligned BAM file, and the reference FASTA sequence.

For each microsatellite locus, the algorithm retrieves all reads overlapping the target region plus user-defined flanking buffer sequence. Reads are filtered to retain only primary alignments, excluding secondary and unmapped reads. The algorithm extracts the aligned portion of each read spanning the locus and determines the microsatellite boundaries within that sequence by identifying regions flanked by at least two consecutive perfect motif repeats at both ends.

Within the identified microsatellite-containing region, the algorithm decomposes the sequence into a structural representation by distinguishing between perfect motif matches and intervening non-motif sequences. For each motif-length window, the algorithm checks for exact matches to the motif or its reverse complement. Consecutive perfect motifs are collapsed into a single block annotated by repeat count and motif sequence (e.g., “5GGAA”). Non-motif intervening sequences are further classified: sequences ≤4 bp or consisting of a single nucleotide (homopolymers) are annotated as-is, single nucleotide variants of the motif are annotated with the variant sequence, and longer complex sequences are recursively decomposed by minimizing edit distance to candidate repeat units or retained as unique sequences if no clear pattern is identified. The resulting structural representation is a concatenated string describing the full allele composition (e.g., “3GGAA_1GGAT_2GGAA”).

For each locus, reads with identical structural representations are grouped together, and the read support for each unique allele structure is quantified. The algorithm computes allele frequency as the proportion of total read support at that locus. Alleles are called using a two-tier frequency threshold approach: if any allele exceeds a user-defined homozygous threshold (default 0.8), only alleles above this threshold are retained; otherwise, all alleles meeting a lower heterozygous threshold (default 0.20) are retained.

This approach accounts for both homozygous and heterozygous loci while filtering low-frequency artifacts.

The genotype module employs parallel processing to analyze chromosomes concurrently. Final genotyping results are output in a tab-separated format where each row represents a single microsatellite locus and includes genomic coordinates, reference allele structure, alternate allele structures, genotype call, allele lengths, total repeat length, maximum consecutive repeat length, and read support metrics for each called allele.

### visualize module

The vmwhere visualize module generates graphical representations of allele frequency and structure at individual microsatellite loci. Input consists of the tab-separated output from vmwhere genotype containing all samples of interest, a unique region identifier, and a minimum allele frequency for filtering.

Allele structure is displayed as a horizontal stacked bar plot in which each bar represents a unique allele and colored blocks indicate sequence composition. Allele frequency at the locus is represented by gray bars positioned to the left of each allele structure. This visualization enables rapid assessment of allelic diversity, structural variation, and relative abundance of microsatellite variants within a cohort.

### Simulated benchmark

#### Simulated loci and read generation

Error-free long-reads were simulated at 30 tetrameric microsatellite loci identified using the vmwhere find module applied to the T2T-CHM13v2 reference genome. For each locus, two distinct alleles were simulated to capture both repeat length variation and sequence-level interruptions. Alleles differed in motif repeat number as well as in the presence and pattern of interrupting sequence motifs within the microsatellite sequence. All microsatellites contained a minimum of one four consecutive repeat segments per the reference defined sequence. Microsatellite bounds were defined as the first and last occurrence of the motif repeat where 2 or more repeats occur within 20 base pairs of a core defined microsatellite.

Simulated reads were generated to a total depth of 30× per locus, with 15× coverage per allele. Each read spanned the full microsatellite allele sequence and included 200 bp of flanking sequence on each side of the repeat region. Reads were written to FASTQ format with uniformly high base quality scores to represent error-free long-read data.

#### Read alignment and BAM processing

The FASTQ containing simulated reads was aligned to the T2T-CHM13v2 reference genome using minimap2(v2.30)(Li 2018, 2021). Resulting alignments were converted to BAM format, sorted and indexed using samtools(v1.23)(Li et al. 2009). Chromosome names were converted to standard UCSC nomenclature from ref seq nomenclature. The same sorted and indexed BAM file was used as input for all microsatellite genotyping tools included in the benchmark.

#### Tool inputs and execution

All tools were provided with identical aligned read data and microsatellite locus coordinates corresponding to the simulated regions. Because tools differed in their expected coordinate conventions (e.g., 0-based versus 1-based indexing), and flanking buffer sizes, locus coordinate files were adjusted as required to fit expected tool input requirements.

The following long-read microsatellite genotyping tools and versions were evaluated. Unless otherwise noted, all tools were executed using their recommended default settings.

vmwhere (v0.1.0), using the genotype module with default parameters

Straglr (v1.5.4), run with tetramer-specific settings (minimum and maximum STR length set to 4 bp), genotype inference enabled based on allele size, a minimum of two supporting reads per allele, and a maximum of three allele clusters per locus

LongTR (v1.2.0), run with default parameters

TRGT (v4.0.0), run in genotype mode with a flanking length of 50 bp

ATaRVa (v0.3.0), run with sequence decomposition enabled

#### Accuracy assessment

Tool outputs were compared to the known simulated ground truth on a per-locus, per-allele basis. Inferred alleles were matched to the two simulated alleles using a minimum absolute difference assignment, whereby predicted alleles were paired to ground-truth alleles to minimize the absolute difference in allele length (in base pairs).

Accuracy was evaluated for multiple genotyping features where available, including allele length (base pairs), repeat length, maximum consecutive repeat length, and motif density. Exact-match accuracy and tolerance-based accuracy were calculated relative to the simulated truth. Wherein a tool made a homozygous call at a microsatellite, accuracy was compared against the closest simulated allele as if the simulated loci was homozygous for that allele.

### 1000 Genomes Project Oxford Nanopore sequencing data

Whole-genome Oxford Nanopore Technology (ONT) sequencing data for the first 100 samples from the 1000 Genomes Project ONT dataset were obtained as unaligned BAM files from the publicly available AWS S3 repository (https://s3.amazonaws.com/1000g-ont/index.html).

Unaligned BAM files were converted to FASTQ format using samtools. FASTQ files were then aligned to the T2T-CHM13v2 human reference genome using minimap2 with parameters optimized for ONT long-read data. Resulting alignments were converted to BAM format, sorted, and indexed using samtools. The resulting sorted and indexed BAM files were used for all downstream microsatellite genotyping and visualization analyses. Sample ancestry annotations were obtained from the 1000 Genomes Project metadata and were used to stratify population-level analyses.

### Comparison of vmwhere find with Repeat Masker and Tandem Repeats Finder

Annotations of repetitive regions generated by RepeatMasker (RM) and Tandem Repeats Finder (TRF) for the T2T-CHM13v2 reference genome were obtained from the UCSC Genome Browser in February 2025. These annotations were converted to BED format where necessary and intersected with the set of tetrameric microsatellite loci identified by vmwhere find using bedtools intersect. Microsatellite loci identified by vmwhere that did not overlap any RepeatMasker or TRF annotations were classified as uniquely detected. Repeat length annotations for all loci were determined directly from the reference sequence using vmwhere.

### Population Scale Analysis

The vmwhere find module was applied to the T2T-CHM13v2 reference genome to identify tetrameric microsatellite loci. Loci containing a minimum of four consecutive repeats were retained and repeat occurrences within 100 bp were merged into a single microsatellite locus. For population-scale analyses, the top ten most frequent tetrameric repeat motifs were selected: AAAC, ATTC, AATG, AAAT, ACAT, AGGG, AAAG, GGAA, AGAT, and ATGG.

Microsatellite genotyping across all samples was performed using the vmwhere genotype module with default algorithm parameters. By default, allele sequences were clustered based on sequence similarity, with alleles within an edit distance of four grouped together. Within each cluster, the allele supported by the highest number of reads was retained as the representative allele. Alleles were required to meet a minimum allele frequency threshold of 0.2, and loci with ≥80% support for a single allele were classified as homozygous. No assumptions of diploidy were imposed. These default settings were applied uniformly across all loci and samples.

As part of genotyping, vmwhere performed sequence decomposition of each allele into contiguous segments based on changes in sequence composition across the repeat sequence. Segments were defined such that any change in repeat sequence, regardless of whether the sequence was canonical or non-canonical, initiated a new segment. As a result, alleles could contain multiple segments alternating between canonical and non-canonical repeat motifs. The number of segments per allele was defined as the total count of these distinct sequence blocks across the full allele sequence.

Genotyping results from all samples and all loci corresponding to the top ten motifs were concatenated for downstream population-scale analyses. For each microsatellite locus, allele counts and frequencies were calculated across the cohort. Singleton alleles—defined as alleles supported by a single read structure at a given locus—were excluded from population-level analyses to reduce spurious allelic calls.

In addition, vmwhere calculated motif density for each allele, defined as the fraction of the allele sequence composed of the canonical tetrameric motif, excluding interrupting sequence variants identified during decomposition.

The variability of each tetrameric sequence was summarized using heterozygosity, computed as 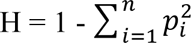, where *p_i_* is the frequency of the *i*-th allele and *n* is the total number of alleles at a locus. Heterozygosity was summarized as a function of maximum consecutive repeat length, with mean heterozygosity calculated across all loci sharing the same maximum consecutive repeat length.

### Oxford Nanopore Technology Whole Genome Sequencing

We performed Oxford Nanopore Technology (ONT) whole genome sequencing on four Ewing sarcoma cell lines.

#### DNA extraction and shearing

DNA was extracted from cell lines using DNeasy Blood & Tissue Kit (Qiagen) following manufacturer’s instructions. Extracted DNA was sheared using a 26G needle for a total of 7 passes to obtain a fragment size distribution for optimal nanopore sequencing throughput. Further size selection was performed using 0.4 volumes of Ampure XP Beads (Beckman Coulter) for cell lines. DNA was quantified using the Qubit fluorometer with the Qubit dsDNA Quantification, High Sensitivity Assay Kit (ThermoFisher Scientific).

#### Library preparation and sequencing

Ligation-based library preparation of native DNA was performed using the following ONT library preparation SQK-LSK 114. SQK-LSK 114 was modified in several ways, including 1) the exclusion of 1 µL DNA CS (DCS) and DCS volume was made up with NFW, 2) volume of 200μL Long Fragment Buffer (LFB) used instead of 250μL LFB, and 3) reduction in the final room temperature incubation time of 5 min instead of the listed 10 min. Samples were sequenced singly on FLO-PRO114M flow cells for 72 h. Samples were sequenced on a PromethION 2 Solo (P2) machine. Reads were base-called (and de-multiplexed, if applicable) using Dorado (v0.5.1-0.6.0[MA2.1]) in super-accurate duplex mode. Base-called reads were aligned to the T2T human reference genome using minimap2.

### ChIP seq, ATAC-seq, and RNA-seq data processing and analysis

#### EWS-FLI1 ChIP-seq

EWS-FLI1 ChIP-seq data were obtained from GEO accession GSE176400. Raw FASTQ files were processed using the nf-core/chipseq pipeline (version 2.1.0) implemented in the Nextflow framework (Ewels et al. 2020; H. Patel et al. 2024) with default parameters.

For read-based quantification, GGAA microsatellites located within 500 bp of one another were merged into a single locus, and the characteristics of the longer microsatellite were retained based on reference-defined repeat length. EWS-FLI1 binding was quantified at these merged loci from each sample BAM file using featureCounts, and signal was normalized to counts per million (CPM).

For visualization of aggregate binding profiles, EWS-FLI1 signal at microsatellites was summarized using the computeMatrix function from deepTools (v3.5.6)(Ramírez et al. 2014), with 500 bp standard microsatellite coordinates bed file and CPM-scaled BigWig files as input. Median signal profiles were visualized using plotProfile with standard error as plot type.

#### ATAC-seq

ATAC-seq data were processed using the nf-core/atacseq pipeline (version 2.1.2) within the Nextflow framework (Ewels et al. 2020; H. Patel et al. 2023) with default parameters. BAM files from biological replicates were merged prior to downstream analyses.

For chromatin accessibility quantification, microsatellites within 500 bp were merged, and the characteristics of the longer microsatellite were retained based on reference-defined repeat length. Accessibility was quantified using featureCounts on merged BAM files and normalized to counts per million (CPM).

#### RNA-seq

Processed microarray data were obtained from Supplementary Table 3A of (Orth et al. 2022) which provides a gene-by-sample expression matrix with values normalized as transcripts per million (TPM). For each Ewing sarcoma cell line (A673, SK-ES-1, MHH-ES-1, and SK-N-MC), TPM values were averaged across replicates to generate a single expression value per gene per cell line.

GGAA microsatellites were annotated with all genes located within 100 kb of each locus. Gene expression values were merged with microsatellite–gene annotations based on gene name. For microsatellite–gene pairs in which the nearby gene was not present in the expression matrix, expression values were recorded as missing, and such entries were omitted from downstream visualization.

### Modeling chromatin accessibility as a function of repeat length

To assess the relationship between microsatellite length and chromatin accessibility, a piecewise linear model was fit using the lm function in R (v4.4.0). The model was

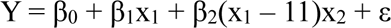

where Y represents log2(CPM +1), x_1_ is the repeat length of microsatellite i, x_2_ is an interaction term for the length of the microsatellite i above 11 repeats and ε is the independent error term.

### Differential accessibility and microsatellite length variation

Differential accessibility testing was performed using DESeq2 (Love, Huber, and Anders 2014) in a one vs rest framework. Log_2_ fold change estimates were shrunk using apeglm (Zhu, Ibrahim, and Love 2019) and loci with an adjusted p-value less than 0.05 were considered differentially accessible.

Differentially accessible peaks were annotated with overlapping microsatellites using bedtools intersect, requiring a minimum overlap of 1 bp. Peaks were permitted to overlap multiple microsatellites and could therefore be represented more than once. Differential accessibility results were merged with microsatellite genotyping information, including maximum consecutive repeat length.

Sample-specific microsatellite expansions and contractions were defined using a one-versus-rest framework. A microsatellite was classified as expanded or contracted if the difference between the largest and smallest repeat length across cell lines was ≥ 2 consecutive repeats.

### Allele specific chromatin accessibility and EWS-FLI1 binding at GGAA microsatellites

#### Haplotype resolved alignment and read processing

A haplotype-resolved *de novo* assembly was generated from long-read whole-genome Oxford Nanopore sequencing data of Ewing cell line A673 using hifiasm (v0.25.0) (Cheng et al. 2021, 2022, 2024). Contig names were verified to be unique, and the two haplotypes were concatenated into a single FASTA file to serve as a combined haplotype reference.

A673 ATAC-seq and EWS-FLI1 ChIP-seq FASTQ reads were aligned to the combined haplotype reference using bwa-mem2 (Vasimuddin et al. 2019). Resulting alignments were sorted by read name using samtools, and samtools fixmate was applied to ensure proper pairing of paired-end ATAC-seq reads (ChIP seq reads are single end, so this step is skipped). PCR duplicates were marked and removed using samtools markdup, and BAM files were indexed with samtools index.

Reads were filtered to retain only primary, high-confidence alignments by excluding secondary, supplementary, unmapped, and quality-control–failed reads, and by requiring a minimum mapping quality of 20. To ensure allele-specific assignment, multi-mapped reads were removed by retaining only reads with an NH tag equal to 1. For ATAC-seq data, filtered BAM files from biological replicates were merged, sorted, and indexed prior to downstream analyses. ChIP seq for EWS-FLI1 and IP did not contain replicates.

#### Allele specific quantification and visualization

To quantify allele-specific chromatin accessibility and EWS-FLI1 binding at GGAA microsatellites, microsatellite loci were first identified directly in the haplotype-resolved *de novo* assembly. The vmwhere find module was applied to the combined haplotype FASTA generated by hifiasm using default parameters (minimum of four consecutive GGAA repeats, with repeat occurrences within 100 bp merged into a single microsatellite). For each GGAA microsatellite, the center of the repeat array was determined, and a fixed window extending ±250 bp from the center was defined for read quantification.

Allele-specific read counts were obtained from the merged replicate, combined haplotype BAM files using featureCounts (Liao, Smyth, and Shi 2014). Because both the microsatellite coordinates and the aligned reads were represented in haplotype-specific contig space, read counts could be assigned independently to each haplotype. For ChIP-seq data, EWS-FLI1 binding signal was normalized using the corresponding input (IP) control reads.

To associate haplotype-resolved microsatellites identified in the *de novo* assembly with genotyped GGAA microsatellites defined in the T2T-CHM13 coordinate space, microsatellite coordinates were lifted over to the T2T reference genome. Lifted-over haplotype-derived microsatellites were then mapped to reference GGAA microsatellite loci using bedtools map (Quinlan and Hall 2010). Repeat length and allele structure for each locus were assigned based on the corresponding vmwhere genotype results.

Microsatellite loci that were homozygous or had insufficient read support (fewer than six ATAC-seq reads across alleles) were excluded from downstream allele-specific analyses. This filtering resulted in a final set of 512 heterozygous GGAA microsatellite loci. Allele-specific signal was summarized as the log₂ ratio of read counts aligned to the longer allele relative to the shorter allele at each microsatellite locus.

To generate haplotype-specific signal tracks for visualization, reads aligned to the combined haplotype reference were separated into haplotype-specific BAM files based on contig naming conventions (haplotype 1 contigs prefixed with h1tg and haplotype 2 contigs prefixed with h2tg). Haplotype-specific alignments were lifted over to the T2T-CHM13 reference genome using a chain file generated from assembly-to-assembly alignment with minimap2 and paf2chain (v0.1.1), followed by coordinate conversion with CrossMap (v0.7.3) (Zhao et al. 2014). Haplotype-specific BAM files were converted to counts-per-million (CPM) normalized BigWig files using deepTools. BigWig files were used exclusively for visualization and were not used for quantitative analyses.

### Figure generation and statistical analysis

All statistical analysis was done using R (v4.4.0). Plots were made using ggplot2 (Wickham 2016). Panels for figures were edited and combined using Adobe illustrator (v2025).

## Acknowledgements

We thank all members of the Ian Davis and Alex Rubinsteyn labs at UNC Chapel Hill for their helpful conversations, and insightful feedback throughout this project. A. McCauley Massie was supported by NIH T32CA244125-05 and the Pediatric Scientist Development Program funded by the Warren Alpert Foundation. IJD is supported by R01CA276663, P30CA016086, the Hyundai Hope on Wheels Foundation, the Wide Open Charitable Foundation of Raleigh, and the Clay on the Green Foundation.

Author contributions are as follows. SKP performed conceptualization, software, formal analysis, visualization, writing - original draft, writing – review & editing. AMM did data curation, investigation, resources, writing - original draft, writing – review & editing. AR did supervision, writing - original draft, writing – review & editing. JRW did software, resources, writing - original draft, writing – review & editing. IJD did conceptualization, supervision, funding acquisition, writing - original draft, writing – review & editing.

