## Supplemental Figures for "Long-read analysis of tetrameric microsatellites with vmwhere supports GGAA repeat length–dependent chromatin state association in Ewing sarcoma"

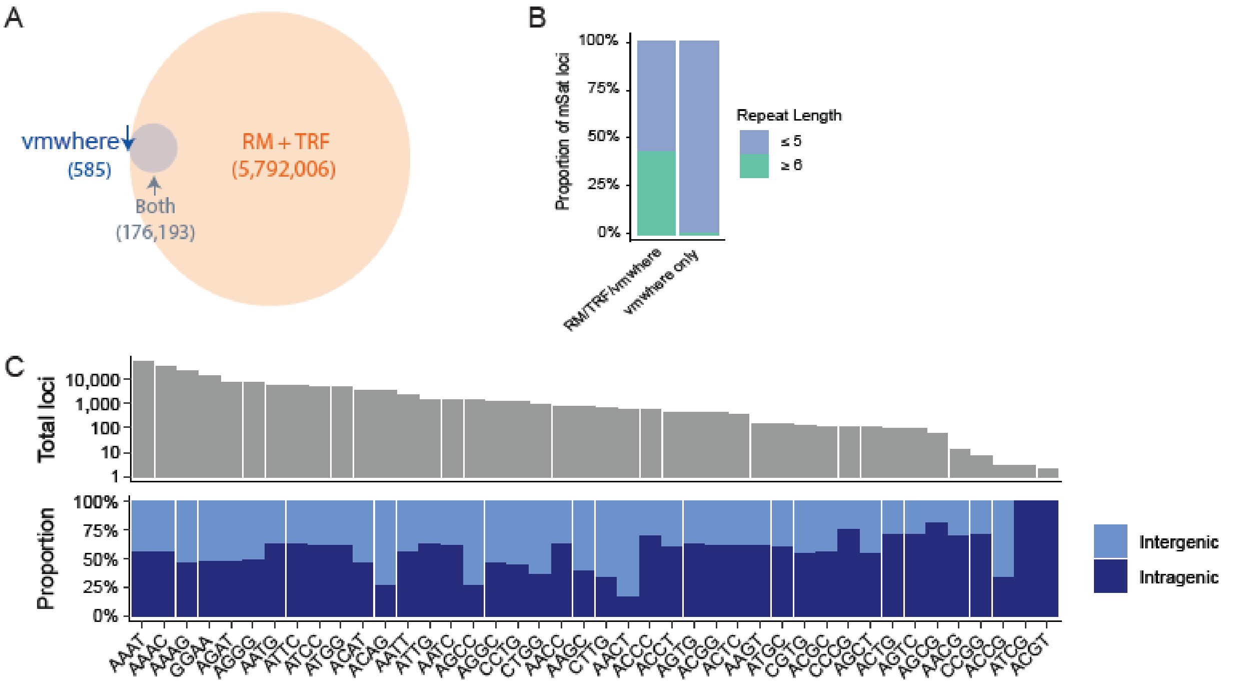


**Supplementary Figure 1. Comparison of microsatellites loci identified via vmwhere find compared to repeat masker and tandem repeats finder**

(A) Overlap between referenced defined coordinates of microsatellite loci using vmwhere and all repeats from Repeat Masker and Tandem Repeats Finder. (B) Microsatellite repeat length based on reference sequence for vmwhere only and shared microsatellite loci. Lengths were determined via vmwhere find. (C) Prevalence of unique tetrameric microsatellite loci in the T2T reference genome (top) and proportion of loci for each motif that are inter or intragenic (bottom).

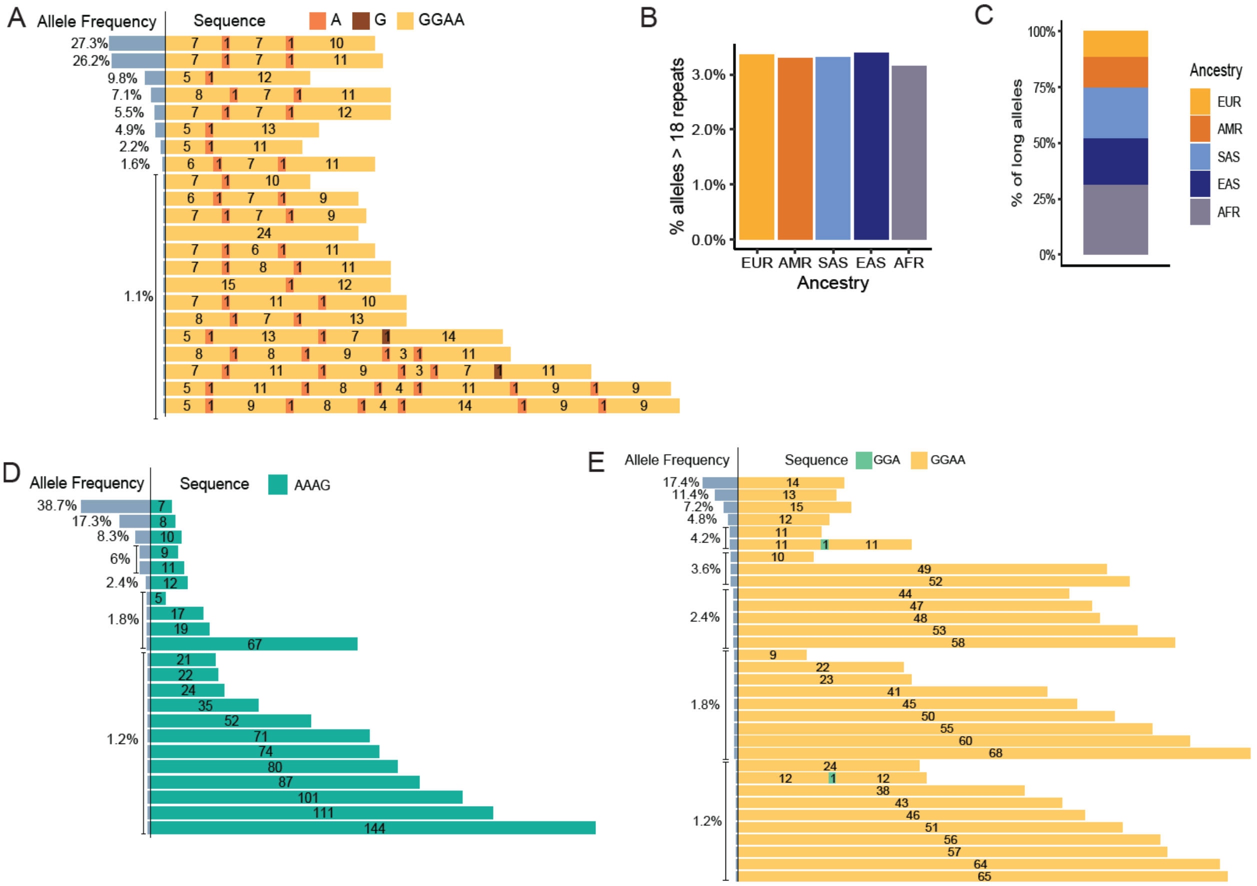


**Supplementary Figure 2. Sequence and ancestry associated allele variation in diverse population of long-read sequenced individuals**

(A) GGAA microsatellite loci exhibits primarily interrupted alleles with several discontinuous segments (B) Percent of alleles, stratified by ancestry, with repeat length greater than 18 (72 base pairs). (C) All long alleles stratified by ancestry, showing percent of long allele associated with each ancestry. (D) AAAG microsatellite loci showing distinct population alleles and frequencies, with all alleles showing uninterrupted repeat lengths. 144 repeat length allele is the longest consecutive repeat length for any AAAG allele. Shorter allele repeat lengths tend to occur at higher frequency (D) GGAA microsatellite loci showing distinct population alleles and frequencies. 68 consecutive repeat length allele is the longest across any GGAA microsatellite loci. Shorter repeat lengths tend to occur at higher frequencies.

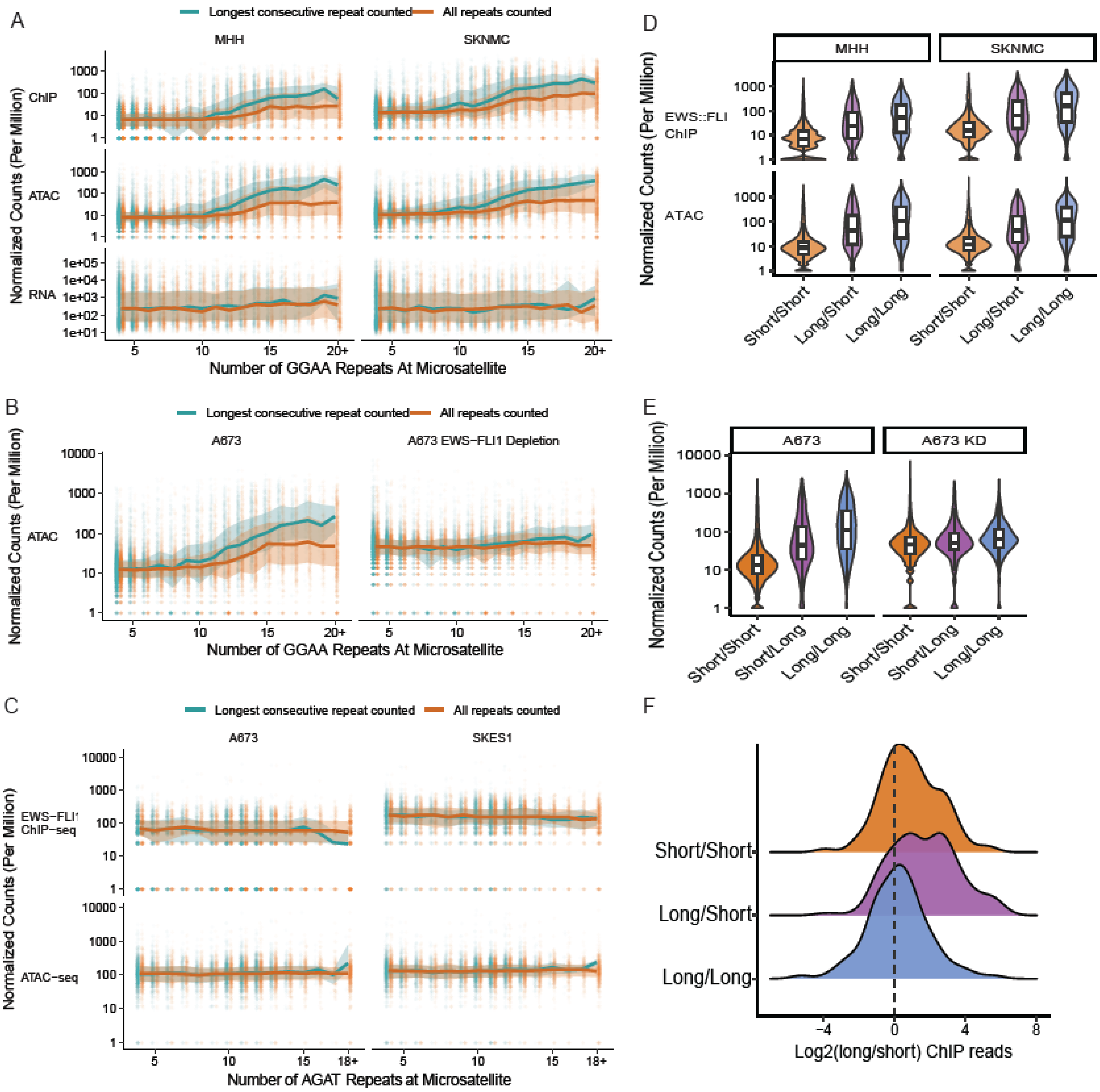


### Supplementary Figure 3. Repeat length associated chromatin state is GGAA specific and EWS-FLI1 dependent in Ewing sarcoma cell lines

(A) Relationship between GGAA repeat length of the longest allele, as determined by vmwhere, and normalized signal (counts per million) at each microsatellite in two Ewing sarcoma cell lines, MHH-ES-1 (left) and SK-N-MC (right). Shown are EWS-FLI1 ChIP–seq (top), ATAC–seq (middle), and RNA–seq (bottom). Normalized signal was quantified once per sequencing modality per microsatellite and is stratified by either maximum consecutive repeat length or total repeat length characteristic at the concordant microsatellite. Line represents the median signal, with shading showing the interquartile (Q1 and Q3) range. (B) Normalized counts for chromatin accessibility at microsatellite loci, stratified by control (left) or EWS-FLI1 knock down condition (right). (C) Normalized signal at AGAT microsatellites, stratified by consecutive repeat length or total repeat length, in A673 (left) and SK-ES-1 (right). (D)

EWS-FLI1 binding and chromatin accessibility at GGAA microsatellites stratified by haplotype, with alleles classified as short (≤11) or long (≥12) based on maximum consecutive repeat length in MHH-ES-1 and SK-N-MC cell lines. (E) Chromatin accessibility at GGAA microsatellites stratified by haplotype in A673 cells with and without EWS-FLI1 knockdown. (F) Distribution of allele specific EWS-FLI1 binding ratios between the two alleles at each microsatellite, quantified as log2 reads long vs short allele at heterozygous microsatellite loci in A673.

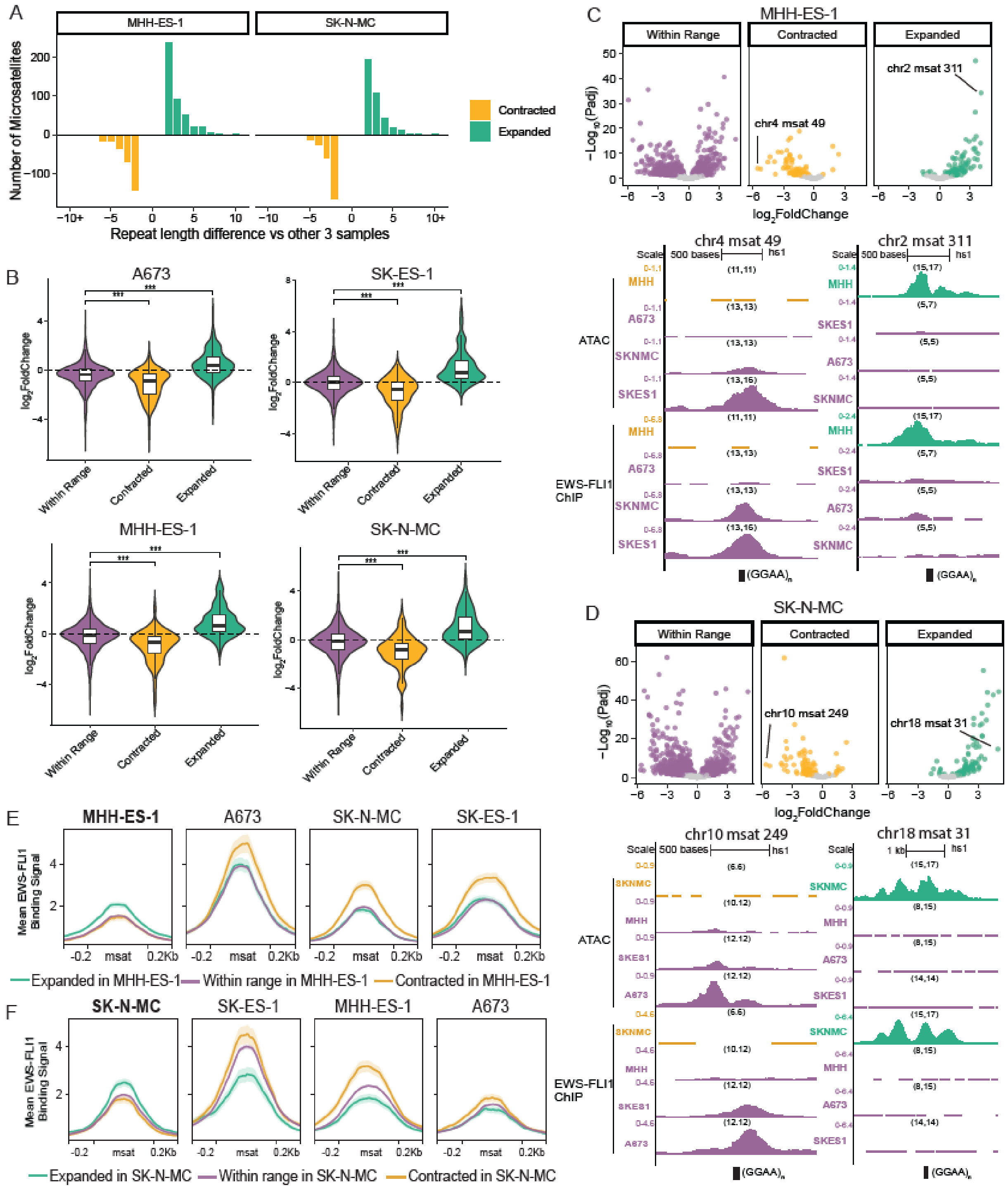


**Supplementary Figure 4. Chromatin accessibility and EWS-FLI1 binding variation mirror repeat length variation in Ewing sarcoma cell lines**

(A) Distribution of consecutive repeat length differences observed among expanded and contracted GGAA microsatellites across cell lines, MHH-ES-1 (left) and SKNMC (right). (B) Violin/box plot per cell line of log2FC chromatin accessibility from one-vs-rest comparison. Microsatellites are stratified via their length variation and log2FC are plotted for each group. Significance determined via Wilcoxon rank-sum test ($p < 0.05$, $p < 0.01$. $p < 0.001$). (C) Volcano plot of differential chromatin accessibility in the MHH-ES-1 cell line, stratified by GGAA microsatellite length change category. The x-axis is log2fold change and the y-axis is -log10(adjusted p-value). Microsatellites within range of other cell lines (left), with allelic support for contraction (middle), or with allelic support for expansion (right) are shown. Points with $p > 0.05$ are shown in grey, and significant loci ($p \leq 0.05$) are colored by microsatellite change category. Example GGAA microsatellite loci in MHH-ES-1 cell line showing the largest decrease in chromatin accessibility associated with repeat contraction (left) and the largest increase associated with repeat expansion (right). ATAC–seq and EWS-FLI1 ChIP–seq signals are shown across all four Ewing sarcoma cell lines. Numbers in parentheses indicate maximum consecutive repeat lengths for the two alleles in each sample at that locus. (D) Volcano plot of differential chromatin accessibility in the SK-N-MC cell line, stratified by GGAA microsatellite length change category, as in panel C. Example GGAA microsatellite loci in SK-N-MC cell line showing the largest decrease in chromatin accessibility associated with repeat contraction (left) and the largest increase associated with repeat expansion (right), displayed as in panel C. Numbers in parentheses indicate maximum consecutive repeat lengths for the two alleles in each sample at that locus. (E) Mean EWS-FLI1 binding signal across GGAA microsatellite loci stratified by expansion status as defined in MHH-ES-1 and shown across all cell lines. (F) Mean EWS-FLI1 binding signal across GGAA microsatellite loci stratified by expansion status as defined in SK-N-MC and shown across all cell lines.
